# Imaging unlabeled axons in the mouse retina by second harmonic generation

**DOI:** 10.1101/2022.05.16.492107

**Authors:** Arafat Meah, Vinessia Boodram, Hyungsik Lim

## Abstract

We describe a novel microscopy for studying the network of axons in the unlabeled fresh wholemount retina. The intrinsic radiation of second harmonic generation (SHG) was utilized to visualize single axons of all major retinal neurons, i.e., photoreceptors, horizontal cells, bipolar cells, amacrine cells, and the retinal ganglion cells. The cell types of SHG+ axons were determined using transgenic GFP/YFP mice. New findings were obtained with retinal SHG imaging: Müller cells do not maintain uniformly polarized microtubules in the processes; SHG+ axons of bipolar cells terminate in the inner plexiform layer (IPL) in a subtype-specific manner; a subset of amacrine cells, presumably the axon-bearing types, emits SHG; and the axon-like neurites of amacrine cells provide a cytoskeletal scaffolding for the IPL stratification. To demonstrate the utility, retinal SHG imaging was applied for testing whether the inner retina is preserved in glaucoma, using DBA/2 mice as a model of glaucoma and DBA/2-*Gpnmb+* as the non-glaucomatous control. It was found that the morphology of the inner retina was largely intact in glaucoma and the pre-synaptic compartments to the retinal ganglion cells were uncompromised. It proves retinal SHG imaging as a promising technology for studying the physiological and diseased retina in 3D.

## 1. Introduction

Imaging the retina’s three-dimensional architecture is crucial for understanding the parallel processing of visual information. The neural circuit has been dissected in the physiological tissue via fluorescent labeling. Green fluorescent protein (GFP) allows targeting specific cells genetically^1-4^. However, due to an expression too low for revealing thin processes, it often requires additional labeling such as Lucifer Yellow or biocytin^5^. Synthetic fluorophores can be microinjected to permit in situ correlation with the neurite’s anatomy during electrophysiological recording, but the low throughput is ill-suited for a large-scale interrogation. Here we describe a new method to image single axon-like processes in the fresh retina without labeling. Endogenous SHG, previously demonstrated for imaging the retinal ganglion cell (RGC) axons^6,7^, can visualize all major neurons in the inner and outer retina.

## 2. Results

### Intrinsic SHG visualizes neurites across the inner and outer retina

SHG can be obtained from the axon because of uniformly polarized microtubules^9,10^. The nonlinear optical radiation is particularly intense from the retinal nerve fiber bundle owing to a large number of RGC axons^6,7^. Although, in principle, SHG should arise also from the axons of non-RGC retinal cells, the signal was expected to be much weaker. Using a setup collecting the forward emission at a half of the excitation wavelength (Fig. 1a), we acquired images of the whole retina. The vertical fibers as well as neuropils were visible in the inner and outer retina (Fig. 1b). The nuclear and plexiform layers could be distinguished for 3D reconstruction (Fig. 1c) and segmentation (Fig. 1d). Major retinal neurons were recognizable, including the photoreceptors and horizontal cells in the outer retina. The emission was not autofluorescence (Suppl. Fig. 1a), but exhibited the characteristics of SHG due to microtubules: It vanished when the retina was fixed with paraformaldehyde (Suppl. Fig. 1b)^11^. The signal was excitable from 700 to 1250 nm, unusually broad for fluorescence. Also, it responded to a microtubule-depolymerizing agent (nocodazole) with the intensity gradually decreasing after the treatment. These results indicated that the new optical contrast is SHG due to axonal microtubules.

**Figure 1.**
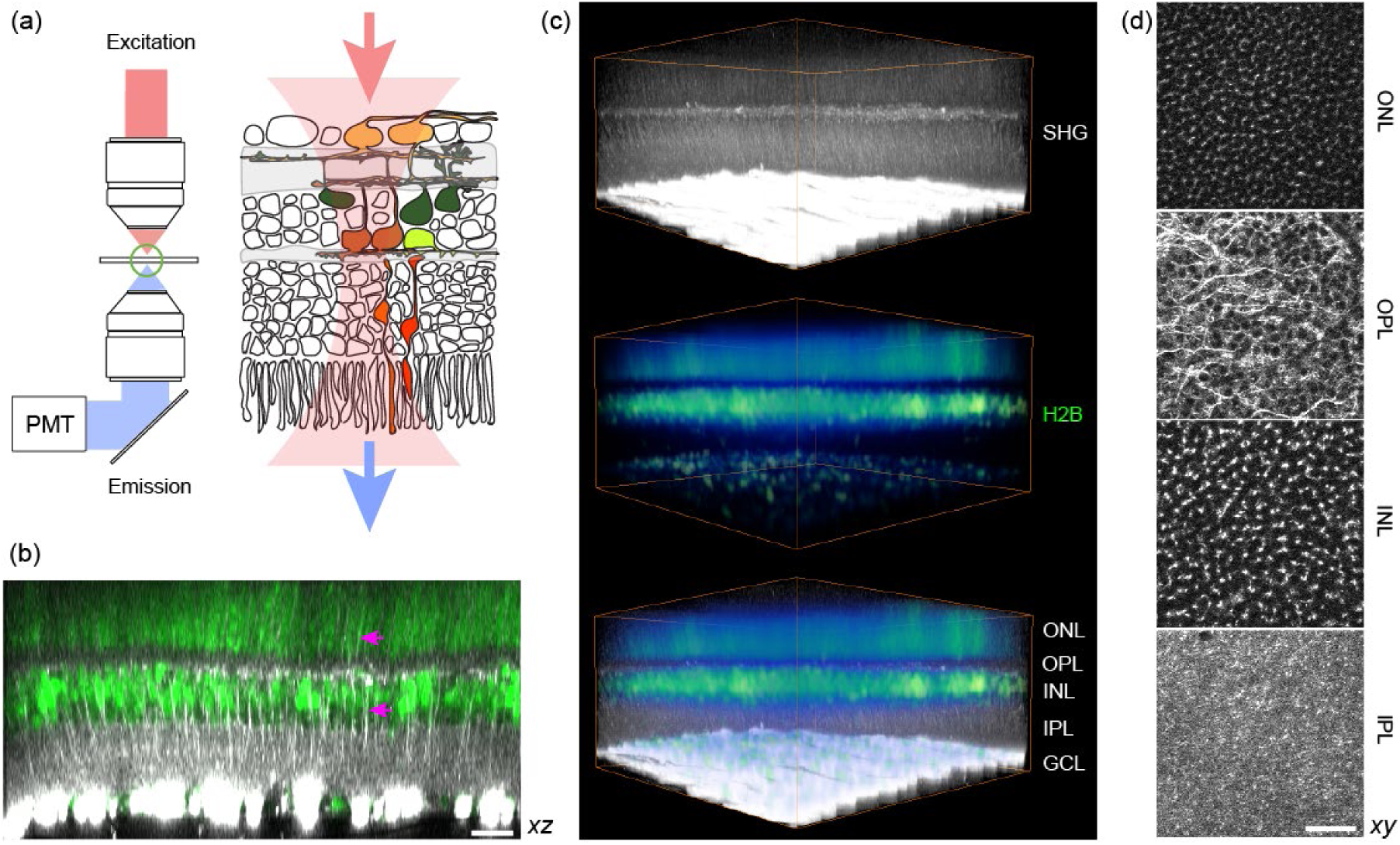
Visualizing the fresh wholemount retina by SHG. (a) Experimental configuration with transmission detection of SHG. (b) Multiphoton imaging of the CAG-H2B-EGFP retina^8^ (an axial projection). The plexiform and nuclear layers are segregated by SHG (gray scale) and EGFP (green), respectively. Vertical SHG processes are also visible (arrows). (c) Volumetric rendering (371×371×186 μm^3^). (d) The segmented layers (lateral projection): ONL, the outer nuclear layer; OPL, the outer plexiform layer; INL, the inner nuclear layer; and IPL, the inner plexiform layer. Scale bars, 30 μm.

### Vertical SHG+ fibers of the inner retina are bipolar cell axons

It seemed that the vertical SHG+ fibers of the inner retina were the axons of bipolar cells. However, Müller cells could not be ruled out completely as certain glial cells are also known to contain polarized microtubules in their processes, e.g., oligodendrocyte^12^. To resolve the cellular identity, the vertical SHG+ fibers were examined in transgenic mice expressing GFP or YFP under the neuro-and glia-specific promoters, i.e., GUS^13^, Thy1 (16 line)^14^, and GFAP^15^. SHG and GFP/YFP were excited simultaneously with a single 915-nm beam. The co-localization was detected in GUS and Thy1-16, but not in GFAP retinas (Fig. 2a). Thus, it was determined that the vertical SHG processes belong to bipolar cells and Müller cells do not polarize microtubules in their processes. Comparing SHG versus GFP/YFP in the same axons revealed that the axon terminals lack SHG signals. Intriguingly, the bipolar cell axons as seen by SHG exhibited variable terminal depths in the IPL (Fig.2a, arrows), which was also observable in vertical slices with a higher resolution (Figs. 2b, c).

**Figure 2.**
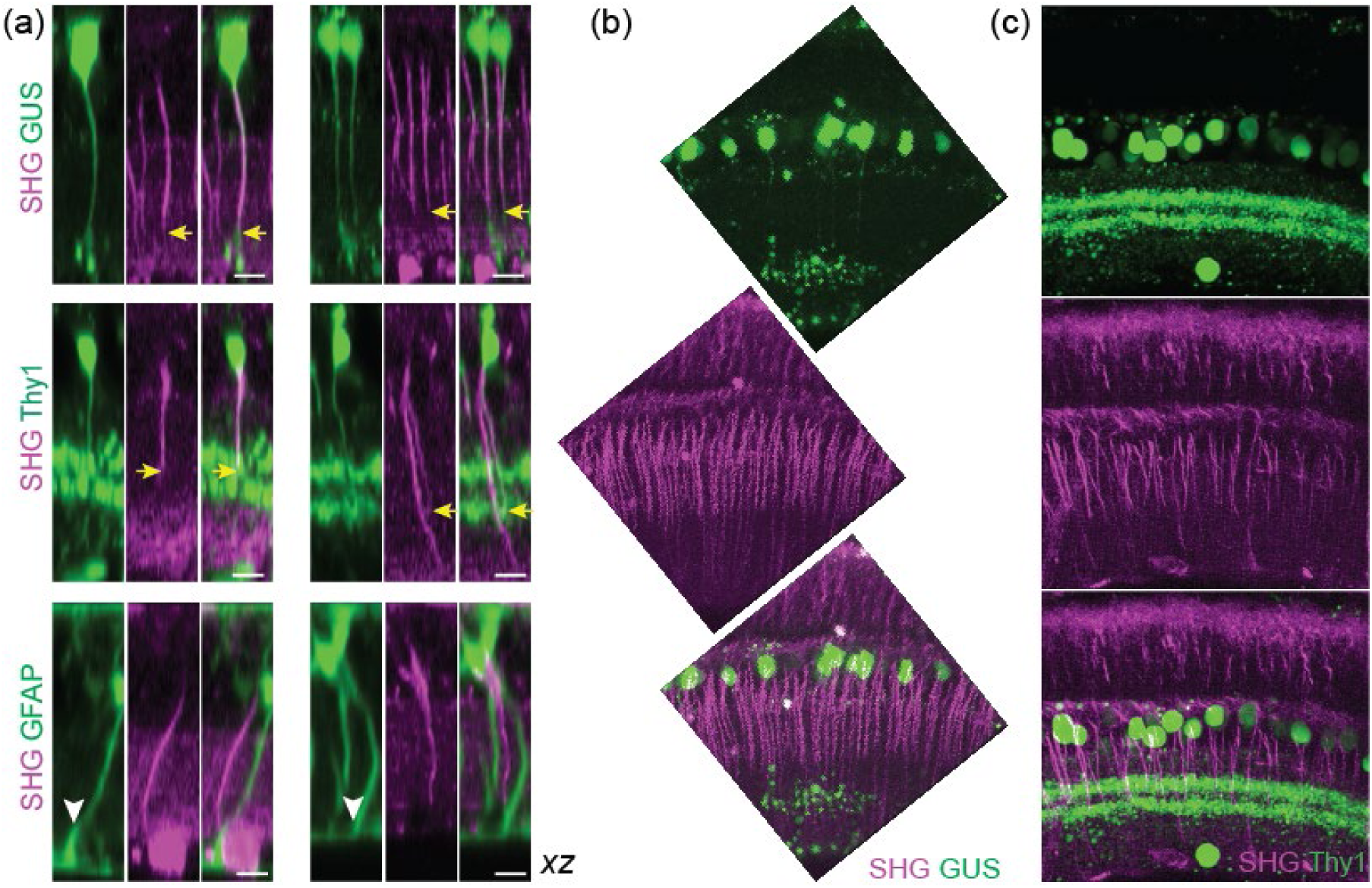
Bipolar cell axons visualized by SHG. (a) Co-registration of SHG and GFP/YFP in GUS-, Thy1-, and GFAP-GFP/YFP mice. Overlaps in GUS and Thy1 (arrows), but not in GFAP retinas (arrowheads). Scale bars, 10 μm. (b), (c) Vertical slices of GUS and Thy1 retinas, respectively.

### The depth of SHG axon terminal is a property of bipolar cell subtype

Bipolar cell ramifies to an IPL sublamina to form synapses with specific amacrine cells and RGCs, therefore the depth of axon terminal is a chief anatomical marker of the function^16,17^. We investigated this parameter with SHG imaging. The census of bipolar cell axons was conducted (Fig. 3a). The number of axons increased steadily toward the INL and then decreased inside the INL, as some SHG fibers coalesced into the interstitial space between cell bodies beyond the optical resolution (Fig. 3b, arrowheads). The peak density, which occurred around zero IPL depth (i.e., the interface between the IPL and INL), was approximately 40,000 to 45,000 axons/mm^2^. It was consistent with the previous counts of bipolar cells in the mouse retina (∼40,000 to 50,000 cells/mm^2^), measured by immunohistochemical staining^18,19^ or electron microscopy^20,21^. Detecting thinnest bipolar cell axons can be improved with a high signal-to-background ratio and a lateral resolution of SHG imaging. The derivative of the number of bipolar cell axons with respect to the depth yielded the number of axon terminals (Fig. 3c), which revealed a subdivision with 4-5 maxima of relatively uniform separation, suggestive of the internal structure of IPL (arrowheads, Fig. 3c). Conceivably, the pattern mimicking the IPL subdivision emerged because of the diverse subtypes of bipolar cells. To test this idea, we examined SHG axon terminals in the GUS-GFP retinas in which three subtypes of bipolar cell are distinguishable, i.e., types 4, 7 cone and rod bipolar cell (CBC4, CBS7, and RBC, respectively)^22^. For an objective measurement of axon terminals, single-neurite tracing was performed on SHG z-stacks (Fig. 3d). The GUS+ subpopulation of SHG traces were identified by superimposition with GFP at zero IPL depth (Fig. 3d). The axial distribution of GUS+ SHG axon terminals showed distinct populations of CBC4, CBC7, and RBC (Fig. 3e). Furthermore, the histogram conformed to the mean GFP profile, verifying the accuracy of measurement. The predicted cell subtypes of the GUS+ SHG axons were corroborated with GFP images (Fig. 3f, g); two cells with SHG axons ending in layer 4 and 5 (blue and yellow arrowheads, respectively) showed characteristic axon terminals of cone and rod bipolar cells, respectively. Consequently, SHG-based classification of bipolar cell subtypes was validated. The demonstrated principle is scalable to the entire retina, and the precision is limited by the axial resolution of imaging (∼1.5 μm).

**Figure 3.**
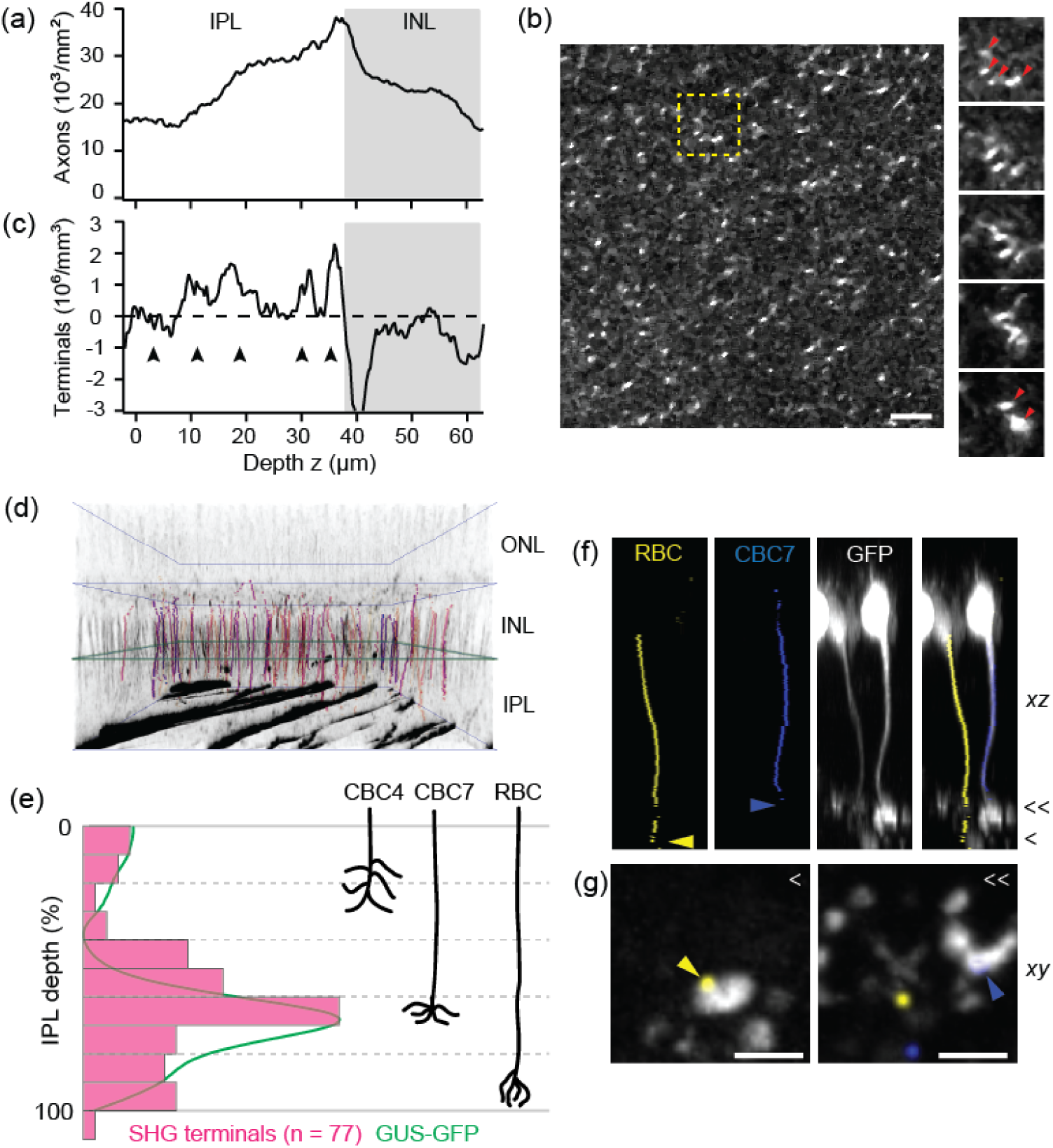
Quantification of the bipolar cell axons. (a) The density of bipolar cell axons. (b) A lateral section of the IPL. Right, fly-through sections corresponding to the dashed box with bipolar cell axons converging into the interstitial space of the INL (arrowheads). Scale bar, 10 μm. (c) The number of bipolar cell axon terminals, estimated by the derivative of (a), exhibiting an internal structure of five layers (arrowheads). (d) 3D rendering of the GUS+ subpopulation of single SHG axons (colored and overlaid with grayscale SHG), identified by the overlap with GFP at zero IPL (green square). (e) The distribution of GUS+ SHG axon terminals (n=77, magenta) and the mean GFP profile (green). Right, the depths of three GUS+ species (CBC4, 7, and RBC) for comparison. (f), (g) GFP+ axon traces overlaid with GFP images, confirming the IPL depth and the characteristic morphology of RBC and CBC7 axon terminals. Scale bars, 5 μm.

### SHG intensity varies significantly along the bipolar cell axons

SHG intensity depends on the amount of uniformly polarized microtubules thus scales with the axon caliber, which is of electrotonic significance but too small to measure by light microscopy. Despite significant variations, the SHG intensity of bipolar cell axon, on average, was higher close to the INL implying the thickening of axons (Fig. 4a). The variations could stem from the subtypes of bipolar cells, e.g., OFF cells have thicker axons than ON cells. However, the SHG intensities of GUS+ axons displayed just as significant variations as total bipolar cell axons at zero IPL depth (Fig. 4b). To disentangle the variability, the SHG intensity was analyzed axially along single axon traces. The individual traces showed little correlations between the SHG intensity of and the IPL depth of the axon terminals. Hence, the cell type was not a major determinant of the SHG intensity. Moreover, the variations of SHG intensity were substantial along the single axons regardless of the subtype, often relative to the choline acetyltransferase (ChAT) bands^3,23-25^ (gray, Fig. 4c). The fluctuating axon diameters are likely to have an implication on the local modulation of visual signals.

**Figure 4.**
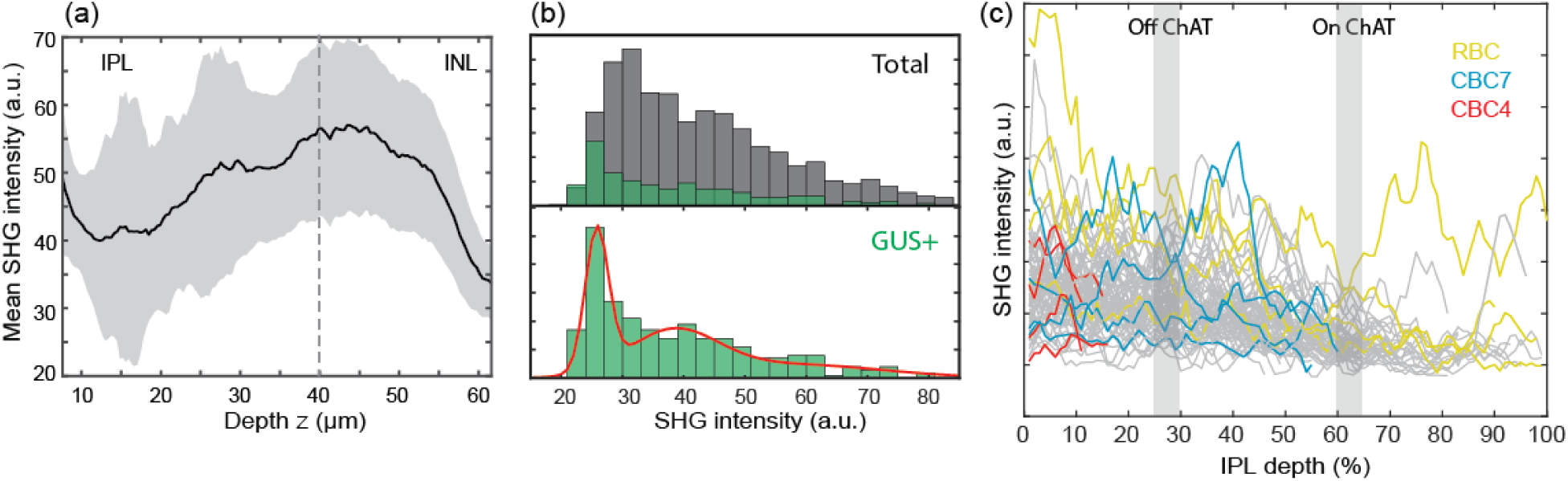
The SHG intensities of individual bipolar cell axons. (a) The mean SHG intensity increases toward the INL. (b) Histograms of the SHG intensity at zero IPL depth of total (gray) and GUS+ (green) bipolar cell axons. Red line, the maximum likelihood fit to Gaussian mixture model which has 3 components. (c) The SHG intensities along the GUS+ axons (n=77).

### Cytoskeletal substrate for the IPL stratification

The IPL consists of 5 (or 10) sublaminae of equal thickness participating in discrete parallel processing^17,26^. SHG imaging of the IPL neuropil routinely revealed a subdivision of 3 strata (Fig. 5a). The axial positions of the gaps coincided approximately with the ChAT bands (Fig. 5a, right), suggesting that the SHG strata aligned with the functional sublayers of sustained ON, transient ON-OFF, and sustained OFF. Furthermore, it indicated that there exists a microtubule scaffolding underlying the IPL stratification, i.e., a cytoskeletal substrate for retinal neurons to establish proper synapses. More refined SHG strata appeared in a shorter range, i.e., ∼5 sublayers over 20 μm (Fig. 5b), implying variable lateral frequencies of SHG+ neurites. To verify the relationship with the IPL sublaminae, SHG strata were analyzed in the GUS-GFP and ChAT-EYFP retinas. The GUS-GFP signal appeared in the fourth of 5 SHG strata (Fig. 5c), while the ChAT-EYFP bands were correctly at the 25 and 62% IPL depths relative to the SHG neuropil (Fig. 5d).

**Figure 5.**
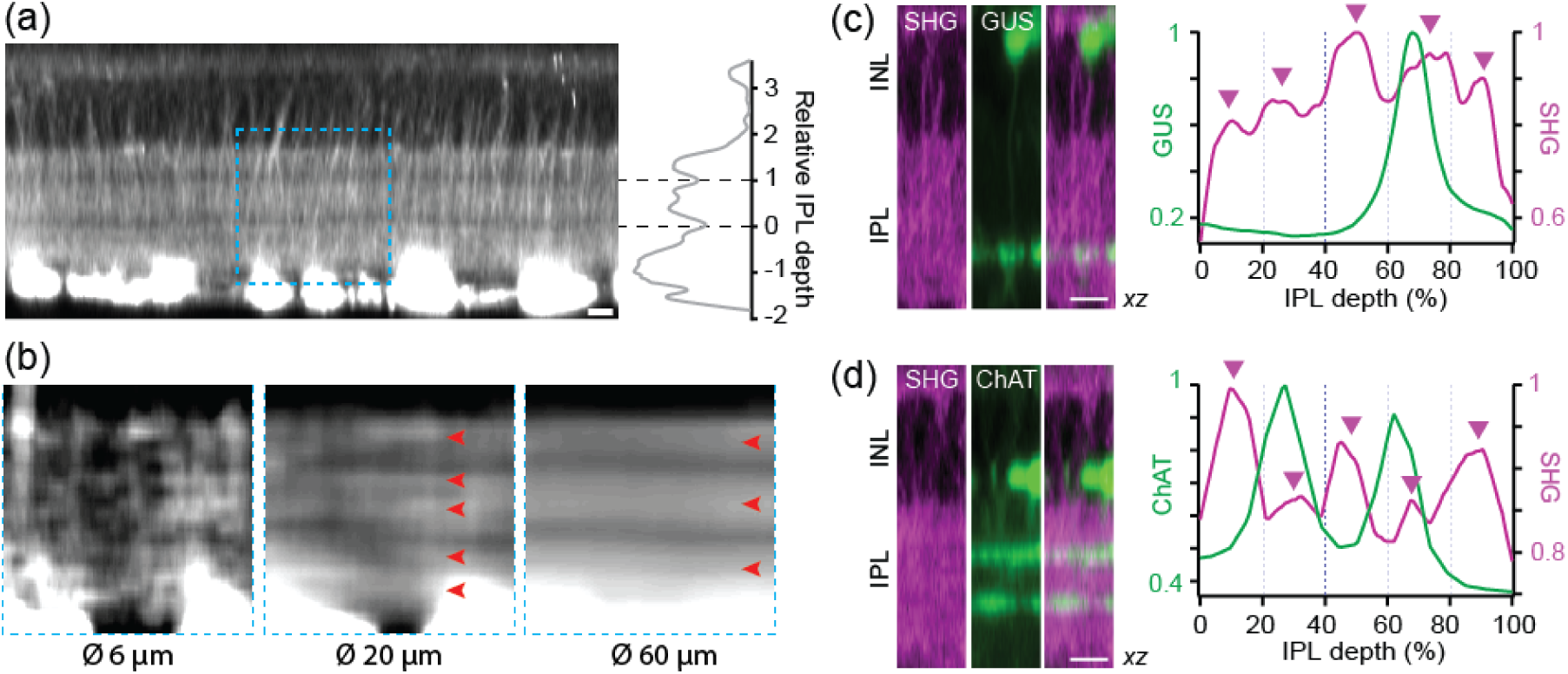
Cytoskeletal substrate underlying the IPL stratification. (a) Two intra-IPL gaps divide SHG strata approximately at the depth of ChAT bands. (b) A region, corresponding to the dashed box in (a), is averaged laterally over 6, 20, and 60 μm. (c), (d) The profiles of SHG versus GFP/YFP in the GUS-GFP and ChAT-EYFP retinas, respectively. The positions of 5 SHG strata are marked with arrowheads. Scale bars, 10 μm.

### Lateral SHG+ processes within the IPL represent a subset of amacrine cells

The molecular origin of the IPL neuropil SHG was determined as microtubules, based on the response to the nocodazole treatment where the intensity of neuropil SHG decreased at the same rate as that of the retinal nerve fibers (Fig. 6a). We asked whether the IPL neuropil SHG was due to specific cell types. The IPL consists of three major neuronal components, i.e., RGC dendrites, amacrine cell neurites, and bipolar cell axon terminals. To find out which is responsible for neuropil SHG, we employed the transgenic retinas of Thy1-YFP (lines H and 16), where the RGC dendrites and amacrine cell neurites are labeled, respectively^14^, and GUS-GFP. SHG and GFP/YFP did not co-localized in GUS-GFP and Thy1-YFP-H (Fig. 6b), excluding bipolar cell axon terminals or RGC dendritic arbors as the source. By contrast, overlaps in Thy1-YFP-16 (arrowheads) suggested that neuropil SHG originates from amacrine cell processes. The observed diffuse SHG in the IPL could result from dense network of sub-resolution neurites.

**Figure 6.**
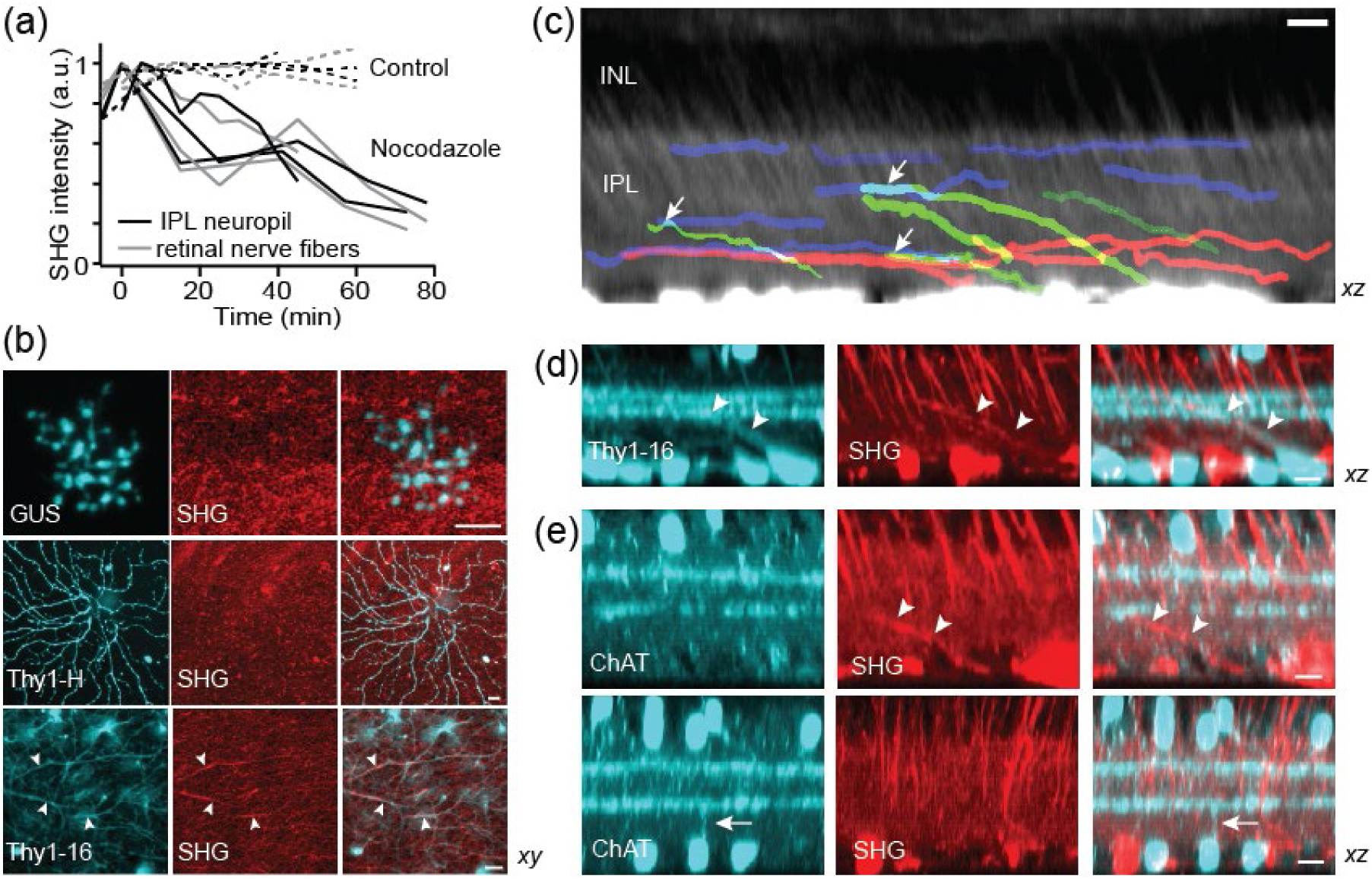
The origin of the IPL neuropil signal. (a) The SHG intensity after the treatment with nocodazole. (b) The co-registration of SHG and GFP/YFP in the GUS, Thy1-YFP-H and -16 retinas. Overlaps only in Thy1-YFP-16 (arrowheads). (c) The lateral SHG+ traces, showing the stratification of mid-(blue), long-range (red), and displaced amacrine cells neurites (green). (d), (e) The identity of SHG+ displaced cell. SHG+ neurite is Thy1+ but not ChAT+ (arrowheads). Conversely, the neurites of ChAT cells are SHG-(arrow). Scale bars, 10 μm.

As the most diverse retinal interneurons, amacrine cells perform a variety of functions in the visual integration, which are dictated by the spatial range of neurites^27-29^. We investigated the lateral field and axial stratification of SHG+ amacrine cells. Single-neurite tracing revealed mostly medium to wide fields (>100 μm) but scarcely narrow fields (Fig. 6c), implying that only a subset of amacrine cells gives rise to SHG. Named for the presumed absence of axons, some amacrine cells are now known to bear axon-like processes (e.g., polyaxonal amacrine cells)^28,30-33^. Plausibly, these axon-bearing amacrine cells are SHG+. In terms of stratification, SHG revealed discrete classes of amacrine cells, i.e., mono-(Fig. 6c, blue and red) and multi-stratified (green). The latter appeared to be displaced amacrine cells whose soma are in the ganglion cell layer. Displaced amacrine cells encompass multiple subtypes whose lateral fields range from narrow to wide^34^. One of the best characterized narrow-field, and also the most numerous, displaced amacrine cells is the starburst amacrine cell (SACs)^23,35^,36. Since SACs are not endowed with axon-like processes, we reasoned that their processes would be SHG-. To test this, we imaged the ChAT-EYFP retinas in which SACs are labeled by EYFP. SHG+ processes of displaced amacrine cells, while Thy1+ (Fig. 6d), were not ChAT+, vice versa (Fig. 6e, arrow and arrowheads). Together with the absence of SHG from the ChAT bands (Fig.5), this result indicated that SAC processes were devoid of uniformly polarized microtubules.

### The broad excitability of SHG aids safer retinal imaging

Since imaging non-RGC cells required an excitation about >5 times higher than for the RGC axons, the risk of photodamage was elevated. There are measures to improve the photosafety. First, the laser beam could be switched off during the fly-back motion of the galvoscanner, i.e., when image is not acquired, reducing the average power that the sample receives by ∼37% with no effect on the image quality. In addition, the excitation could be tuned to a longer near-infrared wavelength, which is in general safer than shorter wavelengths^37,38^. Unlike fluorescence, SHG can be excited at any wavelength since it does not hinge on molecular absorption and similar image qualities were obtained from 700 to 1250 nm (Fig. 7a). To assay the photosafety of SHG imaging, regions in the fresh retina were irradiated at 800, 950, and 1150 nm at the same energy density (∼500 J/cm^2^) and then evaluated by autofluorescence and SHG. The photodamages were negligible at 1150 nm compared to 800 nm (Fig. 7b). Also, the SHG intensity from the ganglion cell layers reduced significantly under the continuous illumination at 800 nm, indicating the loss of cytoskeleton, while it was relatively maintained at 950 and 1150 nm (Fig. 7c). Another indicator of photodamage was provided by the swelling of the retina, likely a result of Müller cell gliosis^39^, which depended on the wavelength. The threshold light dosage to initiate the expansion was much higher at 950 nm than at 800 nm (Fig. 7d). The effect was not noticeable at 1150 nm, where almost no visual transduction was evoked, establishing the relative safety of longer excitation wavelengths.

**Figure 7.**
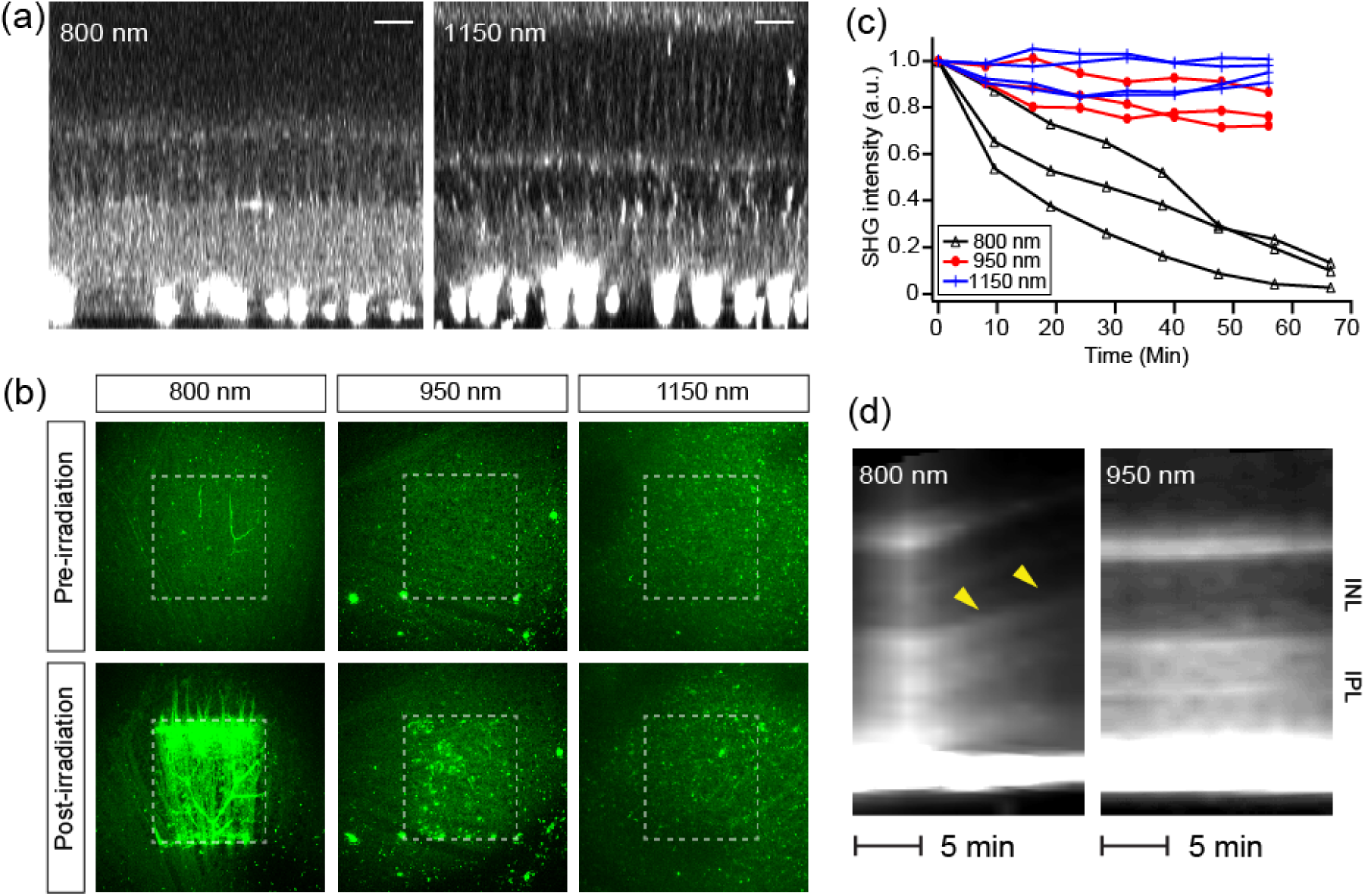
Retinal SHG imaging is tunable to longer wavelengths for safety. (a) SHG imaging at *λ*_*Ex*_=800 and 1150 nm. Scale bars, 20 μm. (b), (c) Light-induced changes evaluated by autofluorescence and SHG, respectively. (d) SHG kymographs of the retina under the continuous illumination. Axial swelling at 800 nm (arrowheads) but not at 950 nm.

### The persistence of the inner retina in glaucoma

To prove the utility, retinal SHG imaging was recruited for asking an outstanding question in glaucoma; namely, whether the pathology is restricted to the RGCs or also involves non-RGC parts of the inner retina. In addition to the RGC dendrites and synapses that degrade in early glaucoma^40-45^, the presynaptic components of amacrine cells and bipolar cell axons might be impaired. The notion has been tested by various modalities in primates and rodents, but the results have been inconsistent^46-49^. We evaluated the inner retina of a mouse model of glaucoma DBA^50,51^ and the non-glaucomatous control DBA-*Gpnmb+*^52^ (n=7 and 4 retinas, respectively, 13-16 months old). Regions around the central retina (0.5 mm from the optic nerve head) were imaged at an excitation wavelength of 950 nm. The SHG images of DBA retinas revealed that, despite substantial loss of the RGC axons, the morphology of the IPL was largely intact (Fig. 8a, b). Also, the SHG intensity in the IPL was comparable to that of DBA-*Gpnmb+*, suggesting that the axon-like neurites of amacrine cells were preserved. Our results complement the previous immunohistochemistry data confirming the structural integrity of amacrine cells in glaucoma^53-56^. Intact amacrine cells may be crucial for glaucoma therapy, not only for preserving the cytoskeletal scaffolding of the IPL but also to maintain amacrine cell-derived signals required for the growth of the RGC axons^57^.

**Figure 8.**
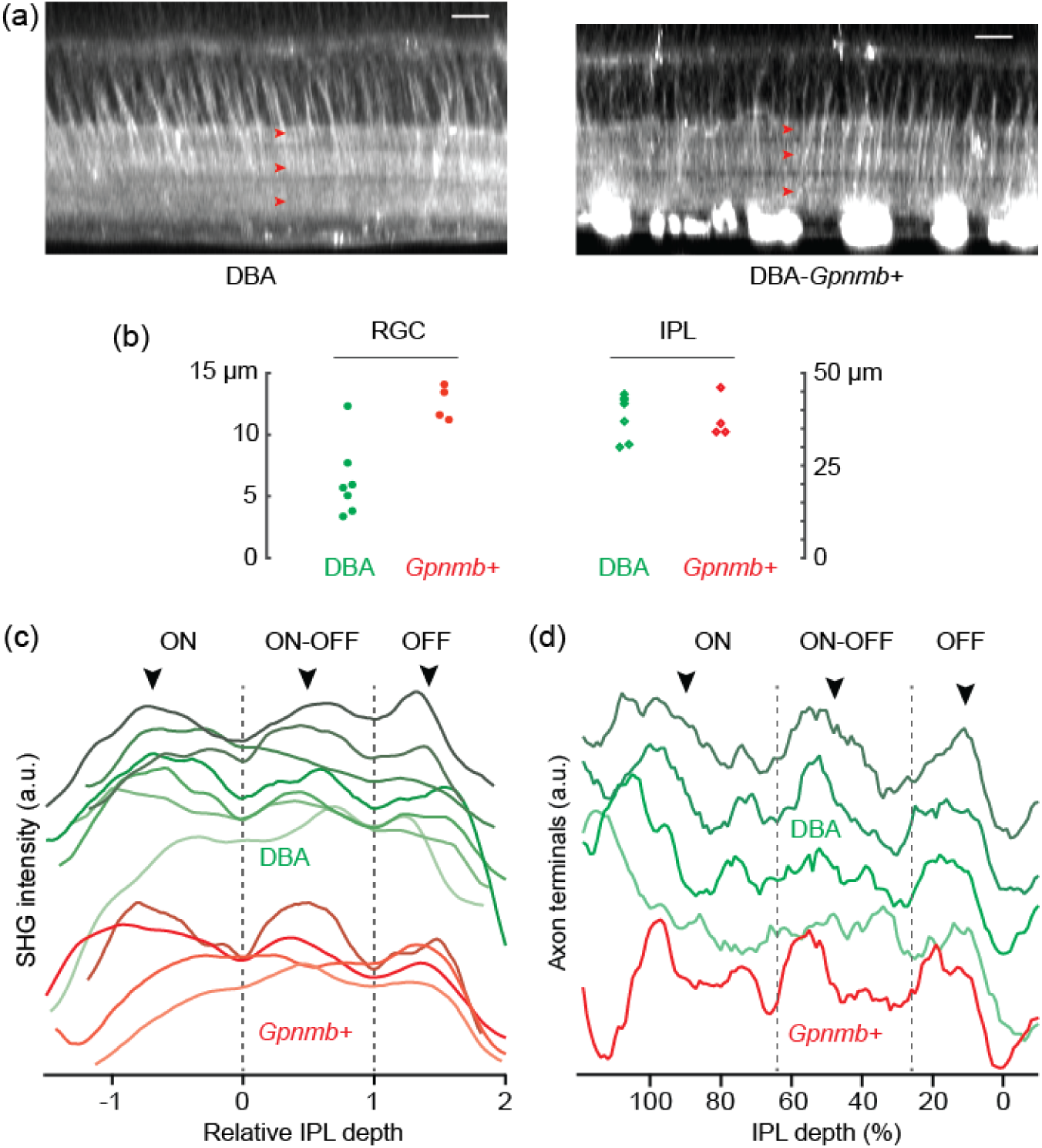
Morphological changes in the glaucomatous inner retina. (a) Representative SHG images of DBA and DBA-*Gpnmb+*. The IPL sublaminae appear normal in both strains (arrowheads). Scale bars, 20 μm. (b) The thicknesses of the RGC axon bundles vs the IPL. (c) The SHG intensity of the IPL neuropil. (d) The density of axon terminals of bipolar cells.

Evidence suggests the progression of glaucoma is circuit-dependent^58,59^; e.g., the ON and OFF pathways exhibit divergent susceptibilities at the level of the RGCs^60-63^. We studied the asymmetry in the larger inner retina by SHG imaging. First, the IPL sublaminae were analyzed for differential degeneration between DBA and DBA-*Gpnmb+*. Three SHG strata maintained the relatively normal thicknesses in DBA (Fig. 8c, arrowheads), indicating the resilience of sustained ON, transient ON-OFF, and sustained OFF layers. Next, the density of bipolar cell axons was compared. The ON and OFF bipolar cells were distinguished by the depth of axon terminals inside the IPL neuropil. The density of axon terminals showed distinct populations of ON, ON-OFF, and OFF cells (Fig. 8d, arrowheads), indicating all the subtypes were largely intact. Taken together, SHG imaging data verified the persistence of the non-RGC axons of the inner retina in glaucoma and no morphological disparity was found between the ON and OFF pathways.

## 3. Discussion

### SHG as an axon-like phenotype

SHG is shown to visualize the axons of all five major classes of retinal neurons. Amacrine cells are the most diverse among them, reflecting the multitude of their functions in the visual processing. With technological advances, new subtypes of amacrine cells have been discovered^1,4^,32,64,65, but the taxonomy may be still incomplete. SHG is obtained only from a certain group of amacrine cells, presumably axon-bearing kinds comprising as much as ∼30% of amacrine cells in the mouse retina^1,30^,32. The axon-like properties of amacrine cell neurites include morphology, spatial range, electrical spiking^33^, and immunoreactivity against phosphorylated neurofilament-H (pNF-H)^66^. Our study raises microtubule polarity as another axonal phenotype, adding a new dimension to the classification of amacrine cells^67^. It opens a set of new questions, e.g., whether all axon-bearing amacrine cells are SHG+; whether the uniform polarity is maintained across the entire length of neurites; and how SHG correlates with other axon-like phenotypes such as pNF-H immunoreactivity.

### Remote interaction between retinal cells in neurodegeneration

Retinal SHG imaging is demonstrated for analyzing the morphology of the inner retina in glaucoma. Determining the precise loci of neurodegeneration is a prerequisite to proper understanding pathogenesis and developing therapeutic strategies. We tested whether the asymmetric susceptibility of ON and OFF RGCs has a remote cause instead of being RGC-autonomous. Considering that the dichotomy initiates in the outer retina^68,69,^ it seemed possible that the intermediate cells connecting the inner and outer retinas might be involved. However, our retinal SHG imaging did not yield any evidence for the hypothesis. The demonstrated ability to survey the large 3D retina without fixation or sectioning can be useful for unraveling the relationship between distant cells in other retinal degeneration, e.g., retinitis pigmentosa^70^, where significant remodeling of the inner retina follows after the degeneration of photoreceptors^71^.

### Comparison to other techniques

SHG imaging can be performed using the same setup as two-photon excited fluorescence microscopy, which is increasingly common in the vision research^35,72-78^. Along with the usual benefits of light microscopy, SHG brings new advantages to retinal imaging. As an intrinsic contrast, it is available from most species from teleost to mammals thus can facilitate comparative studies without transgenic animals. In contrast to other intrinsic contrasts such as reflectance^79-81^ or autofluorescence^74,75^, SHG visualizes the IPL subdivision and the vertical fibers, which are salient features of visual processing. Furthermore, the relatively specific molecular origin makes the data interpretation straightforward.

## 4. Materials and Methods

*Animals*. All procedures were approved by the Hunter College Institutional Animal Care and Use Committee (IACUC). All mice were obtained from The Jackson Laboratory and housed in the animal facility at Hunter College: CAG-H2B-EGFP (B6.Cg-Tg(HIST1H2BB/EGFP)1Pa/J, # 006069), GUS-GFP (Tg(Gnat3-GFP)1Rfm/ChowJ, #026704), Thy1-YFP-H (B6.Cg-Tg(Thy1-YFP)HJrs/J, #003782), Thy1-YFP-16 (B6.Cg-Tg(Thy1-YFP)16Jrs/J, #003709), GFAP-GFP (FVB/N-Tg(GFAPGFP)14Mes/J, #003257), DBA (DBA/2J, #000671), and DBA-*Gpnmb+* (DBA/2J-*Gpnmb+*/SjJ, #007048). ChAT-EYFP mouse was obtained by crossing ChAT-IRES-Cre knock-in (B6.129S-*Chat*^*tm1(cre)Lowl*^/MwarJ (#031661)^82^ with R26R-EYFP reporter mice (B6.129X1-*Gt(ROSA)26Sor*^*tm1(EYFP)Cos*^/J, #006148)^83^.

*Tissue preparation and pharmacology*. The retinal wholemounts were prepared as previously described^6,7^. Animal was euthanized by CO_2_ inhalation and the eye was enucleated. The retinal wholemounts were immersed in the oxygenated Ames’ medium for imaging. It took less than 30 minutes from animal’s euthanasia to the beginning of imaging. For vertical slices, the fresh retinal wholemount was embedded in 6% agar and cut into 250-μm slices with a vibrating blade microtome (Leica VT100 S). For nocodazole treatment, the drug was added to the Ames’ medium at the final concentration of 33 μM.

*SHG microscopy*. SHG microscopy was performed using a setup previously described^6,7^. Short pulses of a 150-fs duration and an 80-MHz repetition rate from a Ti:Sapphire laser tunable from 700 to 1050 nm (Chameleon; Coherent, Inc.) or an optical parametric oscillator tunable from 1050 to 1250 nm (OPO, Angewandte Physik & Elektronik GmbH) were used for excitation. The polarization state of excitation beam was controlled with half-and quarter-waveplates. A Pockels cell (Conoptics 350-80LA) was inserted in the beam path for switching off the laser beam during the fly-back motion of galvoscanner (‘fly-back blanking’), which was driven with an electrical signal synchronized with the galvoscanner. The excitation beam was focused with a water-dipping objective lens (Nikon CFI75 16× 0.8NA or Leica HC FLUOTAR L 25x 0.95NA). The average power was 100-150 mW at the sample. The forward-propagating SHG signal was collected with an objective lens (Olympus UApo340 40× 1.35NA) and detected with a photomultiplier tube (PMT; Hamamatsu H10770PA-40). The SHG channel contained a narrow-bandpass filter (<20-nm bandwidth) with the center wavelength at a half of the excitation wavelength. The pixel dwell time was ∼3 μs. Typically 1-5 frames were acquired at the frame rate of ∼1.5 Hz. The z step of z-stacks images was 2 or 3 μm.

*Image processing and data analysis*. Image processing was done using ImageJ^84^ and MATLAB (MathWorks, Inc.). 3D reconstruction was obtained with Amira (Thermo Scientific). Segmentation was done using Trainable Weka Segmentation plugin^85^. Single-neurite tracing was performed by an automatic procedure modified from single-particle tracking^86,87^ or semi-manually by semiautomatic ridge detection^88,89^.

## Acknowledgements

This work was supported by funding from the National Institute of Health (GM140841).

## Competing interests

None.

---

**Supplementary Figure 1.**
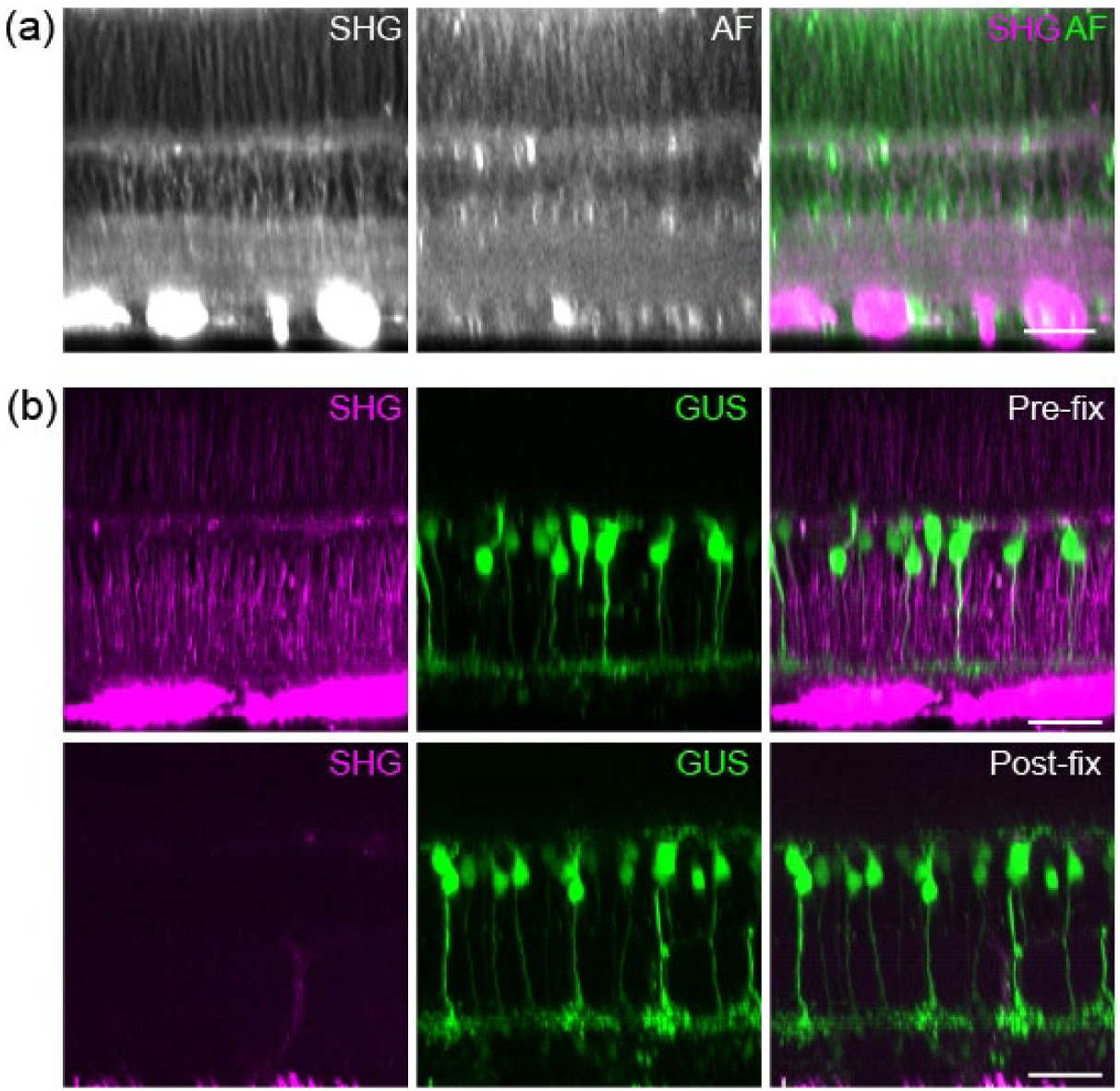
The new optical signal from the retina is SHG. (a) SHG and autofluorescence (AF) from the fresh retinal wholemount. (b) SHG, but not GFP, was lost after paraformaldehyde fixation of the GUS-GFP retina. Scale bars, 30 μm.

## Notes

### Competing Interest Statement

The authors have declared no competing interest.

## References

1 Badea, T. C. & Nathans, J. Quantitative analysis of neuronal morphologies in the mouse retina visualized by using a genetically directed reporter. Journal of Comparative Neurology 480, 331–351, doi:10.1002/cne.20304 (2004).

2 Siegert, S. et al. Genetic address book for retinal cell types. Nat. Neurosci. 12, 1197–U1130, doi:10.1038/nn.2370 (2009).

3 Ivanova, E., Hwang, G. S. & Pan, Z. H. Characterization of transgenic mouse lines expressing Cre recombinase in the retina. Neuroscience 165, 233–243, doi:10.1016/j.neuroscience.2009.10.021 (2010).

4 Zhu, Y. L., Xu, J., Hauswirth, W. W. & DeVries, S. H. Genetically targeted binary labeling of retinal neurons. Journal of Neuroscience 34, 7845–7861, doi:10.1523/jneurosci.2960-13.2014 (2014).

5 Horikawa, K. & Armstrong, W. E. A versatile means of intracellular labeling: injection of biocytin and its detection with avidin conjugates. J. Neurosci. Methods 25, 1–11, doi:10.1016/0165-0270(88)90114-8 (1988).

6 Lim, H. & Danias, J. Label-free morphometry of retinal nerve fiber bundles by second-harmonic-generation microscopy. Optics Letters 37, 2316–2318 (2012).

7 Sharoukhov, D., Bucinca-Cupallari, F. & Lim, H. Microtubule imaging reveals cytoskeletal deficit predisposing the retinal ganglion cell axons to atrophy in DBA/2J. Investigative Ophthalmology & Visual Science 59, 5292–5300 (2018).

8 Hadjantonakis, A. & Papaioannou, V. Dynamic in vivo imaging and cell tracking using a histone fluorescent protein fusion in mice. BMC Biotechnology 4, - (2004).

9 Baas, P. W., Deitch, J. S., Black, M. M. & Banker, G. A. Polarity orientation of microtubules in hippocampal neurons - Uniformity in the axon and nonuniformity in the dendrite. Proceedings of the National Academy of Sciences of the United States of America 85, 8335–8339 (1988).

10 Dombeck, D. et al. Uniform polarity microtubule assemblies imaged in native brain tissue by second-harmonic generation microscopy. Proceedings of the National Academy of Sciences of the United States of America 100, 7081–7086 (2003).

11 Van Steenbergen, V. et al. Molecular understanding of label-free second harmonic imaging of microtubules. Nature Communications 10, doi:10.1038/s41467-019-11463-8 (2019).

12 Lunn, K. F., Baas, P. W. & Duncan, I. D. Microtubule organization and stability in the oligodendrocyte. Journal of Neuroscience 17, 4921–4932 (1997).

13 Huang, L. Q. et al. G protein subunit G gamma 13 is coexpressed with G alpha o, G beta 3, and G beta 4 in retinal ON bipolar cells. Journal of Comparative Neurology 455, 1–10, doi:10.1002/cne.10396 (2003).

14 Feng, G. P. et al. Imaging neuronal subsets in transgenic mice expressing multiple spectral variants of GFP. Neuron 28, 41–51, doi:10.1016/s0896-6273(00)00084-2 (2000).

15 Zhuo, L. et al. Live astrocytes visualized by green fluorescent protein in transgenic mice. Dev. Biol. 187, 36–42, doi:10.1006/dbio.1997.8601 (1997).

16 Ghosh, K. K., Bujan, S., Haverkamp, S., Feigenspan, A. & Wassle, H. Types of bipolar cells in the mouse retina. Journal of Comparative Neurology 469, 70–82, doi:10.1002/cne.10985 (2004).

17 Euler, T., Haverkamp, S., Schubert, T. & Baden, T. Retinal bipolar cells: Elementary building blocks of vision. Nature Reviews Neuroscience 15, 507–519, doi:10.1038/nrn3783 (2014).

18 Jeon, C. J., Strettoi, E. & Masland, R. H. The major cell populations of the mouse retina. Journal of Neuroscience 18, 8936–8946 (1998).

19 Wässle, H., Puller, C., Muller, F. & Haverkamp, S. Cone contacts, mosaics, and territories of bipolar cells in the mouse retina. Journal of Neuroscience 29, 106–117, doi:10.1523/jneurosci.4442-08.2009 (2009).

20 Helmstaedter, M. et al. Connectomic reconstruction of the inner plexiform layer in the mouse retina. Nature 500, 168-+, doi:10.1038/nature12346 (2013).

21 Behrens, C., Schubert, T., Haverkamp, S., Euler, T. & Berens, P. Connectivity map of bipolar cells and photo receptors in the mouse retina. Elife 5, doi:10.7554/eLife.20041 (2016).

22 Lin, B. & Masland, R. H. Synaptic contacts between an identified type of ON cone bipolar cell and ganglion cells in the mouse retina. Eur. J. Neurosci. 21, 1257–1270, doi:10.1111/j.1460-9568.2005.03967.x (2005).

23 Famiglietti, E. V. ‘Starburst’ amacrine cells and cholinergic neurons: mirror-symmetric ON and OFF amacrine cells of rabbit retina. Brain Research 261, 138–144, doi:10.1016/0006-8993(83)91293-3 (1983).

24 Guiloff, G. D. & Kolb, H. Neurons immunoreactive to choline acetyltransferase in the turtle retina. Vision Research 32, 2023–2030, doi:10.1016/0042-6989(92)90063-o (1992).

25 Haverkamp, S. & Wässle, H. Immunocytochemical analysis of the mouse retina. Journal of Comparative Neurology 424, 1–23 (2000).

26 Ramóny Cajal, S. The structure of the retina.. (Charles C. Thomas, 1892).

27 Kolb, H. Amacrine cells of the mammalian retina: Neurocircuitry and functional roles. Eye 11, 904–923, doi:10.1038/eye.1997.230 (1997).

28 MacNeil, M. A. & Masland, R. H. Extreme diversity among amacrine cells: Implications for function. Neuron 20, 971–982, doi:10.1016/s0896-6273(00)80478-x (1998).

29 Masland, R. H. The tasks of amacrine cells. Visual Neuroscience 29, 3–9, doi:10.1017/s0952523811000344 (2012).

30 Vaney, D. I., Peichl, L. & Boycott, B. B. Neurofibrillar long-range amacrine cells in mammalian retinae. Proceedings of the Royal Society Series B-Biological Sciences 235, 203-+, doi:10.1098/rspb.1988.0072 (1988).

31 Dacey, D. M. Axon-bearing amacrine cells of the macaque monkey retina Journal of Comparative Neurology 284, 275–293, doi:10.1002/cne.902840210 (1989).

32 Lin, B. & Masland, R. H. Populations of wide-field amacrine cells in the mouse retina. Journal of Comparative Neurology 499, 797–809, doi:10.1002/cne.21126 (2006).

33 Greschner, M. et al. A polyaxonal amacrine cell population in the primate retina Journal of Neuroscience 34, 3597–3606, doi:10.1523/jneurosci.3359-13.2014 (2014).

34 Muller, L. P. D., Shelley, J. & Weiler, R. Displaced amacrine cells of the mouse retina. Journal of Comparative Neurology 505, 177–189, doi:10.1002/cne.21487 (2007).

35 Euler, T., Detwiler, P. & Denk, W. Directionally selective calcium signals in dendrites of starburst amacrine cells. Nature 418, 845–852 (2002).

36 Zheng, J. J., Lee, S. & Zhou, Z. J. A developmental switch in the excitability and function of the starburst network in the mammalian retina. Neuron 44, 851–864, doi:10.1016/j.neuron.2004.11.015 (2004).

37 Squirrell, J., Wokosin, D., White, J. & Bavister, B. Long-term two-photon fluorescence imaging of mammalian embryos without compromising viability. Nature Biotechnology 17, 763–767 (1999).

38 Debarre, D., Olivier, N., Supatto, W. & Beaurepaire, E. Mitigating phototoxicity during multiphoton microscopy of live Drosophila embryos in the 1.0-1.2 µm wavelength range. PLoS One 9, doi:10.1371/journal.pone.0104250 (2014).

39 Bringmann, A. et al. Muller cells in the healthy and diseased retina. Progress in Retinal and Eye Research 25, 397–424, doi:10.1016/j.preteyeres.2006.05.003 (2006).

40 Weber, A. J., Kaufman, P. L. & Hubbard, W. C. Morphology of single ganglion cells in the glaucomatous primate retina. Investigative Ophthalmology & Visual Science 39, 2304–2320 (1998).

41 Weber, A. J. & Harman, C. D. Structure-function relations of parasol cells in the normal and glaucomatous primate retina. Investigative Ophthalmology & Visual Science 46, 3197–3207, doi:10.1167/iovs.04-0834 (2005).

42 Williams, P. A. et al. Retinal ganglion cell dendritic atrophy in DBA/2J glaucoma. PLoS One 8, doi:10.1371/journal.pone.0072282 (2013).

43 Berry, R. H., Qu, J., John, S. W. M., Howell, G. R. & Jakobs, T. C. Synapse loss and dendrite remodeling in a mouse model of glaucoma. PLoS One 10, doi:10.1371/journal.pone.0144341 (2015).

44 Liu, M., Duggan, J., Salt, T. E. & Cordeiro, M. F. Dendritic changes in visual pathways in glaucoma and other neurodegenerative conditions. Experimental Eye Research 92, 244–250, doi:10.1016/j.exer.2011.01.014 (2011).

45 Agostinone, J. & Di Polo, A. in New Trends in Basic and Clinical Research of Glaucoma: A Neurodegenerative Disease of the Visual System, Pt A Vol. 220 Progress in Brain Research (eds G. Bagetta & C. Nucci) 199–216 (Elsevier Science Bv, 2015).

46 Raz, D., Perlman, I., Percicot, C. L., Lambrou, G. X. & Ofri, R. Functional damage to inner and outer retinal cells in experimental glaucoma. Investigative Ophthalmology & Visual Science 44, 3675–3684, doi:10.1167/iovs.02-1236 (2003).

47 Choi, S. S. et al. Evidence of outer retinal changes in glaucoma patients as revealed by ultrahigh-resolution in vivo retinal imaging. British Journal of Ophthalmology 95, 131–141, doi:10.1136/bjo.2010.183756 (2011).

48 Vidal-Sanz, M. et al. Understanding glaucomatous damage: Anatomical and functional data from ocular hypertensive rodent retinas. Progress in Retinal and Eye Research 31, 1–27, doi:10.1016/j.preteyeres.2011.08.001 (2012).

49 Park, H. Y. L., Kim, J. H. & Park, C. K. Alterations of the synapse of the inner retinal layers after chronic intraocular pressure elevation in glaucoma animal model. Molecular Brain 7, doi:10.1186/s13041-014-0053-2 (2014).

50 John, S. W. M. et al. Essential iris atrophy, pigment dispersion, and glaucoma in DBA/2J mice. Investigative Ophthalmology & Visual Science 39, 951–962 (1998).

51 Bayer, A. U. et al. Retinal morphology and ERG response in the DBA/2NNia mouse model of angle-closure glaucoma. Invest Ophthalmol Vis Sci 42 (2001).

52 Howell, G. R. et al. Absence of glaucoma in DBA/2J mice homozygous for wild-type versions of Gpnmb and Tyrp1. BMC Genetics 8, 45, doi:10.1186/1471-2156-8-45 (2007).

53 Vickers, J. C. et al. Differential vulnerability of neurochemically identified subpopulations of retinal neurons in a monkey model of glaucoma. Brain Research 680, 23–35, doi:10.1016/0006-8993(95)00211-8 (1995).

54 Jakobs, T. C., Libby, R. T., Ben, Y. X., John, S. W. M. & Masland, R. H. Retinal ganglion cell degeneration is topological but not cell type specific in DBA/2J mice. Journal of Cell Biology 171, 313–325 (2005).

55 Kielczewski, J. L., Pease, M. E. & Quigley, H. A. The effect of experimental glaucoma and optic nerve transection on amacrine cells in the rat retina. Investigative Ophthalmology & Visual Science 46, 3188–3196, doi:10.1167/iovs.05-0321 (2005).

56 Gunn, D. J., Gole, G. A. & Barnett, N. L. Specific amacrine cell changes in an induced mouse model of glaucoma. Clinical and Experimental Ophthalmology 39, 555-+, doi:10.1111/j.1442-9071.2010.02488.x (2011).

57 Goldberg, J. L., Klassen, M. P., Hua, Y. & Barres, B. A. Amacrine-signaled loss of intrinsic axon growth ability by retinal ganglion cells. Science 296, 1860–1864, doi:10.1126/science.1068428 (2002).

58 Bui, B. V., Edmunds, B., Cioffi, G. A. & Fortune, B. The gradient of retinal functional changes during acute intraocular pressure elevation. Investigative Ophthalmology & Visual Science 46, 202–213, doi:10.1167/iovs.04-0421 (2005).

59 Kong, Y. X., Crowston, J. G., Vingrys, A. J., Trounce, I. A. & Bui, B. V. Functional changes in the retina during and after acute intraocular pressure elevation in mice. Investigative Ophthalmology & Visual Science 50, 5732–5740, doi:10.1167/iovs.09-3814 (2009).

60 Della Santina, L., Inman, D. M., Lupien, C. B., Horner, P. J. & Wong, R. O. L. Differential progression of structural and functional alterations in distinct retinal ganglion cell types in a mouse model of glaucoma. Journal of Neuroscience 33, 17444–17457, doi:10.1523/jneurosci.5461-12.2013 (2013).

61 Pang, J. J., Frankfort, B. J., Gross, R. L. & Wu, S. M. Elevated intraocular pressure decreases response sensitivity of inner retinal neurons in experimental glaucoma mice. Proceedings of the National Academy of Sciences of the United States of America 112, 2593–2598, doi:10.1073/pnas.1419921112 (2015).

62 El-Danaf, R. N. & Huberman, A. D. Characteristic patterns of dendritic remodeling in early-stage glaucoma: Evidence from genetically identified retinal ganglion cell types. Journal of Neuroscience 35, 2329–2343, doi:10.1523/jneurosci.1419-14.2015 (2015).

63 Ou, Y., Jo, R. E., Ullian, E. M., Wong, R. O. L. & Della Santina, L. Selective vulnerability of specific retinal ganglion cell types and synapses after transient ocular hypertension. Journal of Neuroscience 36, 9240–9252, doi:10.1523/jneurosci.0940-16.2016 (2016).

64 Macneil, M. A., Heussy, J. K., Dacheux, R. F., Raviola, E. & Masland, R. H. The shapes and numbers of amacrine cells: Matching of photofilled with Golgi-stained cells in the rabbit retina and comparison with other mammalian species. Journal of Comparative Neurology 413, 305–326 (1999).

65 Yan, W. J. et al. Mouse retinal cell atlas: Molecular identification of over sixty amacrine cell types. Journal of Neuroscience 40, 5177–5195, doi:10.1523/jneurosci.0471-20.2020 (2020).

66 Volgyi, B. & Bloomfield, S. A. Axonal neurofilament-H immunolabeling in the rabbit retina. Journal of Comparative Neurology 453, 269–279, doi:10.1002/cne.10392 (2002).

67 Zeng, H. K. & Sanes, J. R. Neuronal cell-type classification: challenges, opportunities and the path forward. Nature Reviews Neuroscience 18, 530–546, doi:10.1038/nrn.2017.85 (2017).

68 Werblin, F. S. & Dowling, J. E. Organization of the retina of the mudpuppy, Necturus maculosus. II. Intracellular recording. J. Neurophysiol. 32, 339-& (1969).

69 Kaneko, A. Physiological and morphological identification of horizontal, bipolar and amacrine cells in goldfish retina. Journal of Physiology-London 207, 623-&, doi:10.1113/jphysiol.1970.sp009084 (1970).

70 Strettoi, E. & Pignatelli, V. Modifications of retinal neurons in a mouse model of retinitis pigmentosa. Proceedings of the National Academy of Sciences of the United States of America 97, 11020–11025, doi:10.1073/pnas.190291097 (2000).

71 Marc, R. E., Jones, B. W., Watt, C. B. & Strettoi, E. Neural remodeling in retinal degeneration. Progress in Retinal and Eye Research 22, 607–655, doi:10.1016/s1350-9462(03)00039-9 (2003).

72 Euler, T. et al. Eyecup scope-optical recordings of light stimulus-evoked fluorescence signals in the retina. Pflugers Arch. 457, 1393–1414, doi:10.1007/s00424-008-0603-5 (2009).

73 Wei, W., Elstrott, J. & Feller, M. B. Two-photon targeted recording of GFP-expressing neurons for light responses and live-cell imaging in the mouse retina. Nature Protocols 5, 1347–1352, doi:10.1038/nprot.2010.106 (2010).

74 Imanishi, Y., Batten, M., Piston, D., Baehr, W. & Palczewski, K. Noninvasive two-photon imaging reveals retinyl ester storage structures in the eye. Journal Of Cell Biology 164, 373–383 (2004).

75 Palczewska, G. et al. Noninvasive multiphoton fluorescence microscopy resolves retinol and retinal condensation products in mouse eyes. Nature Medicine 16, 1444–U1130, doi:10.1038/nm.2260 (2010).

76 Borghuis, B. G. et al. Imaging light responses of targeted neuron populations in the rodent retina Journal of Neuroscience 31, 2855–2867, doi:10.1523/jneurosci.6064-10.2011 (2011).

77 Borghuis, B. G., Marvin, J. S., Looger, L. L. & Demb, J. B. Two-photon imaging of nonlinear glutamate release dynamics at bipolar cell synapses in the mouse retina. Journal of Neuroscience 33, 10972–10985, doi:10.1523/jneurosci.1241-13.2013 (2013).

78 Sharma, R. et al. In vivo two-photon imaging of the mouse retina. Biomedical Optics Express 4, 1285–1293, doi:10.1364/boe.4.001285 (2013).

79 Gloesmann, M. et al. Histologic correlation of pig retina radial stratification with ultrahigh-resolution optical coherence tomography. Investigative Ophthalmology & Visual Science 44, 1696–1703, doi:10.1167/iovs.02-0654 (2003).

80 Srinivasan, V. J. et al. 5522–5528.

81 Bizheva, K. et al. Optophysiology: Depth-resolved probing of retinal physiology with functional ultrahigh-resolution optical coherence tomography. Proceedings of the National Academy of Sciences of the United States of America 103, 5066–5071, doi:10.1073/pnas.0506997103 (2006).

82 Rossi, J. et al. Melanocortin-4 Receptors Expressed by Cholinergic Neurons Regulate Energy Balance and Glucose Homeostasis. Cell Metabolism 13, 195–204, doi:10.1016/j.cmet.2011.01.010 (2011).

83 Srinivas, S. et al. Cre reporter strains produced by targeted insertion of EYFP and ECFP into the ROSA26 locus. BMC Developmental Biology 1, 1–8 (2001).

84 Schneider, C. A., Rasband, W. S. & Eliceiri, K. W. NIH Image to ImageJ: 25 years of image analysis. Nature Methods 9, 671–675, doi:10.1038/nmeth.2089 (2012).

85 Arganda-Carreras, I. et al. Trainable Weka Segmentation: a machine learning tool for microscopy pixel classification. Bioinformatics 33, 2424–2426, doi:10.1093/bioinformatics/btx180 (2017).

86 Crocker, J. C. & Grier, D. G. Methods of digital video microscopy for colloidal studies. Journal of Colloid and Interface Science 179, 298–310 (1996).

87 Chenouard, N. et al. Objective comparison of particle tracking methods. Nature Methods 11, 281–U247, doi:10.1038/nmeth.2808 (2014).

88 Meijering, E. et al. Design and validation of a tool for neurite tracing and analysis in fluorescence microscopy images. Cytometry Part A 58A, 167–176, doi:10.1002/cyto.a.20022 (2004).

89 Longair, M. H., Baker, D. A. & Armstrong, J. D. Simple Neurite Tracer: open source software for reconstruction, visualization and analysis of neuronal processes. Bioinformatics 27, 2453–2454, doi:10.1093/bioinformatics/btr390 (2011).

