## Supplementary Figure 1 for "Imaging unlabeled axons in the mouse retina by second harmonic generation"

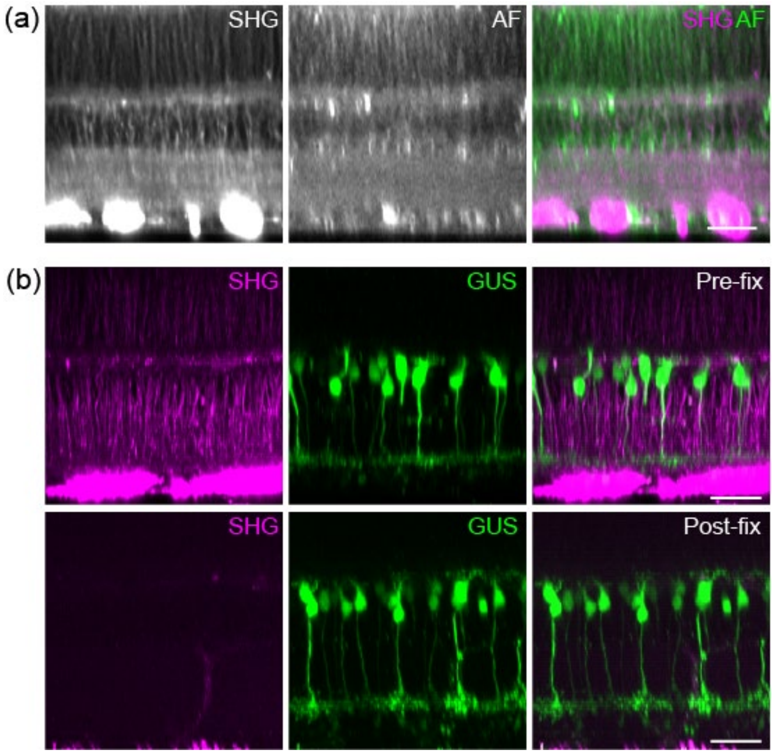


**Supplementary Figure 1**. The new optical signal from the retina is SHG. (a) SHG and autofluorescence (AF) from the fresh retinal wholemount. (b) SHG, but not GFP, was lost after paraformaldehyde fixation of the GUS-GFP retina. Scale bars, 30 μm.
